# Towards a unified theory of efficient, predictive and sparse coding

**DOI:** 10.1101/152660

**Authors:** Matthew Chalk, Olivier Marre, Gašper Tkačik

## Abstract

A central goal in theoretical neuroscience is to predict the response properties of sensory neurons from first principles. Several theories have been proposed to this end. “Efficient coding” posits that neural circuits maximise information encoded about their inputs. “Sparse coding” posits that individual neurons respond selectively to specific, rarely occurring, features. Finally, “predictive coding” posits that neurons preferentially encode stimuli that are useful for making predictions. Except in special cases, it is unclear how these theories relate to each other, or what is expected if different coding objectives are combined. To address this question, we developed a unified framework that encompasses these previous theories and extends to new regimes, such as sparse predictive coding. We explore cases when different coding objectives exert conflicting or synergistic effects on neural response properties. We show that predictive coding can lead neurons to either correlate or decorrelate their inputs, depending on presented stimuli, while (at low-noise) efficient coding always predicts decorrelation. We compare predictive versus sparse coding of natural movies, showing that the two theories predict qualitatively different neural responses to visual motion. Our approach promises a way to explain the observed diversity of sensory neural responses, as due to a multiplicity of functional goals performed by different cell types and/or circuits.

---

Sensory neural circuits perform a myriad of computations, which allow us to make sense of, and interact with, our environment. For example, neurons in the primary visual cortex encode information about local edges in an image [1], while neurons in higher-level areas encode more complex features, such as textures or faces [2, 3]. A central aim of sensory neuroscience is to develop a mathematical theory to explain the purpose and nature of such computations, and, ultimately, predict neural responses to stimuli from first principles.

Several theories have been proposed about the function that sensory systems have evolved to perform. The efficient coding hypothesis posits that sensory circuits transmit maximal information about their inputs, given internal constraints, such as metabolic costs and/or noise [4, 5, 6, 7]. Alternatively, the sparse coding hypothesis posits that individual neurons respond selectively to specific, rarely occurring, features in the environment [8, 9, 10]. Finally, the more recent predictive coding hypothesis^1^ posits that sensory neurons transmit maximal information about stimuli that are predictive about the future while discarding non-predictive information [11, 12].

One may ask which, if any, of these objectives are fulfilled by sensory neural circuits. This question is all the more important given that, in many cases, different coding objectives appear to directly conflict with each other. For example, a classic result of efficient coding in the low-noise regime is that neurons should temporally decorrelate their inputs and preferentially encode fast stimulus features [13, 14, 15, 16]. In contrast, predictive coding favours the extraction of temporally-correlated, slow features [17, 18]. Likewise, sparse coding requires that neurons respond selectively to a single, preferred stimulus feature. It is unclear if this is compatible with predictive coding, which requires neurons to respond to stimuli as quickly as possible.

While a large body of theoretical work exists on efficient and sparse coding (reviewed in [19, 20]), there is little work on how neurons could optimally encode stimuli that are pre dictive about the future (with the exception of [21, 17]); in short, the general implications of predictive coding for neural circuits are still unknown. We also do not understand how different coding objectives relate to each other, or what happens when they are combined.

Here, we incorporate the three existing theories—sparse coding, efficient coding, and predictive coding—into a unified framework. In this framework, a small set of optimisation parameters determines the functional goals and constraints faced by sensory neurons. Previous theories correspond to specific values of these optimisation parameters. As a result, we can investigate the conditions under which different coding objectives, such as encoding predictive information versus maximising efficiency, have conflicting or synergistic effects on neural responses. Further, we can explore qualitatively new coding regimes, such as neural codes that are both predictive and sparse. We end by hypothesizing that the observed diversity of sensory neural responses spans the space of coding tradeoffs accessible by varying the parameters of our new theory.

## A unified framework for predictive and efficient coding

We consider a stimulus, *y*_−∞:*t*_ ≡ (…, *y_t−_*_1_, *y_t_*), giving rise to a sensory input, *x_t_ = y_t_* + *n_t_,* where *n_t_* represents input noise. We look for the optimal neural code, *p*(*r_t_|x*_−∞:*t*_), such that neural responses *r* within a temporal window of length *τ* encode maximal information about the stimulus at time lag Δ, given a fixed amount of information encoded about past inputs, *C* (Fig a).

**Fig. 1.**
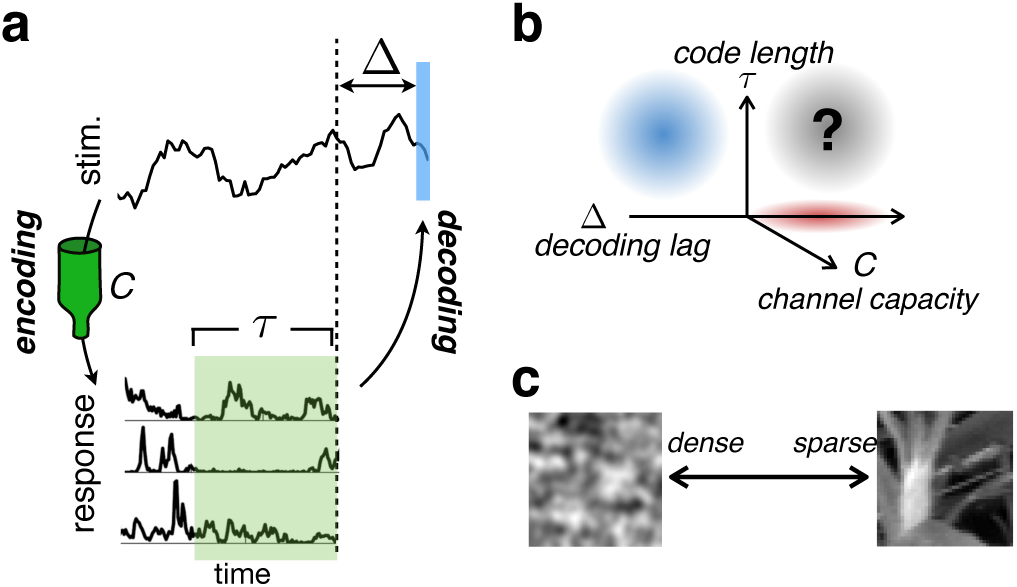
Schematic of modeling framework. **(a)** A stimulus (above) elicits a response in a population of neurons (below). We look for optimal codes, where the responses within a time window of length *τ* maximise information encoded about the stimulus at lag Δ, subject to a constraint on the information about past inputs, C. **(b)** For a given stimulus, the optimal code depends on three parameters: *τ*, Δ, and *C*. Previous work on efficient temporal coding looked at *τ* > 0, and Δ < 0 (blue shade). Previous work on predictive coding looked at Δ > 0 and *τ* ~ 0 (red shade,. Our theory is valid in all regimes, but we focus in particular on Δ > 0 & *τ* > 0 (Mack shade), **(c)** We further explore how optimalcodes change when there is a sparse latent structure in the stimulus (natural image patch, right) vs when there is none filtered gaussian noise, left).

This problem can be formalised using the information bottleneck (IB) framework [22], in which one seeks a code, *p*(*r_t_*|*x*–_∞:*t*_), that maximises the objective function:

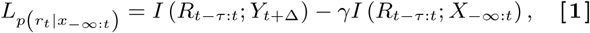

where the first term is the mutual information between responses *R*_*t−τ*:*t*_ and stimulus *Y_t_*_+Δ_, to be maximised, and the second term denotes the mutual information between *R*_*t−τ*:*t*_ and *X*_−∞:*t*_, to be constrained (which we call the channel capacity, *C*). A constant, *γ*, determines the strength of this constraint, and thus, the tradeoff between coding fidelity and compression. When *γ* = 0 the optimal solution is to encode the input perfectly; when *γ* = 1 the optimal solution is to encode zero information about the input.

In general, it is impossible to exactly maximise the objective function in Eq (1). We previously presented an approximate method that instead maximizes the information about the stimulus which can be recovered from neural responses using a specific type of decoder (e.g., a linear decoder); formally, this amounts to a variational approximation that maximizes a lower bound on *L* [23], as described in SI Section 1. To recover efficient coding in the limit where *X* ≈ *Y*, we replaced R*_t_*_−τ:*t*_ in the second term of Eq (1) with the instantaneous response, *R_t_*. With this modification, maximising *L* for Δ < 0 is equivalent to minimising the redundancy in the responses, as in efficient coding (see Table 1 and SI Section 1).

**Table 1.**
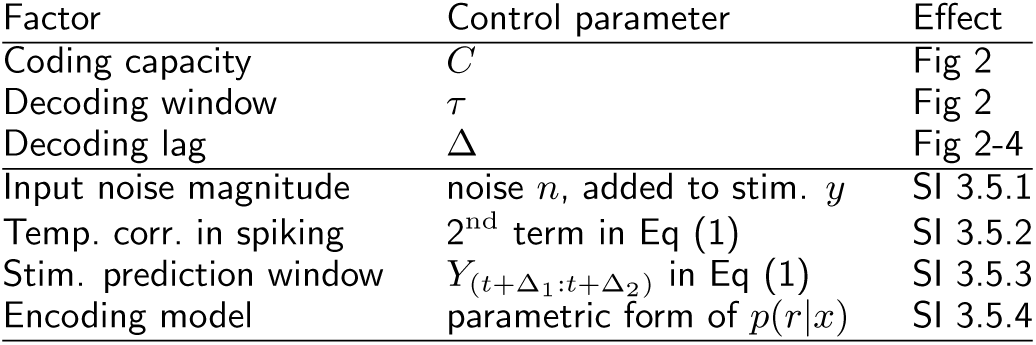
List of factors determining the optimal neural code. The first three factors are explored in detail in the main text.

Equation (1) clearly shows that the optimal coding strategy depends on three factors: the *decoding lag*, Δ, the *code length*, *τ*, and the *channel capacity, C* (determined by 7). Previous theories of neural coding correspond to specific regions within the three-dimensional parameter space spanned by Δ, *τ*, and *C* (Fig b). For example, efficient coding investigated how, at low noise, neurons transmit maximal information about past inputs (Δ < 0) by minimising temporal redundancy in their responses [13, 14]. This strategy is optimal when the stimulus can be read-out by integrating neural responses over time, i.e., when *τ* > 0 (blue region in Fig b). In contrast, predictive coding (Δ > 0) looked exclusively at near-instantaneous codes, where *τ* ~ 0 (red region in Fig b)^2^ [12, 21, 17]. Below, we investigate the relation between these previous works and focus on the (previously unexplored) case of neural codes that are both predictive (Δ > 0) and temporal (*τ* > 0; grey region in Fig b). To specialize our theory to the biologically-relevant case, we further investigate predictive coding of natural stimuli. A hallmark of natural stimuli is their sparse structure [19, 20, 8, 24]: stimulus fragments can be constructed from a set of primitive features (e.g., image contours), each of which occurs rarely (Fig c). By incorporating sparsity into our information-theoretic framework, we explore the relationship between sparse and predictive coding.

## Results

### Dependence of neural code on coding objectives

Our initial goal was to understand the influence of different coding objectives in the simplest scenario, where a single neuron linearly encodes a 1-d input. In this model, the neural response at time *t* is:

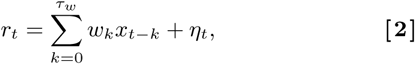

where 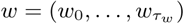 are the linear coding weights and *η_t_* is a gaussian noise with unit variance.

With stimuli that have gaussian statistics, the objective function takes a very simple form:

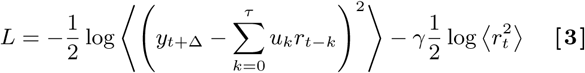

where *u* = (*u*_0_, …, *u*_*τ*_) are the optimal linear read-out weights used to reconstruct the stimulus at time *t* + Δ from the responses between *t* − Δ and *t*. Thus, the optimal code is the one that minimises the mean-squared reconstruction error at lag Δ, constrained by the variance of the neural response (relative to the noise variance).

Initially, we investigated “instantaneous” predictive coding, where *τ* = 0, so that the stimulus at time *t* + Δ is estimated from the instantaneous neural response at time *t* (Fig 2a). We considered three different stimulus types, shown in Fig 2b. With a “Markov” stimulus, whose future trajectory depended on the current state, *y_t_,* only (Fig 2b, top panel; see SI Section 2.1), to predict the stimulus at a future time, *y_t_*_+Δ_, neurons only needed to encode the current state *y_t_*. Thus, when *τ* = 0, we observed the trivial solution where *r_t_* ∝ *y_t_*, irrespective of the decoding lag, Δ (Fig 2c-d, top panels).

**Fig. 2.**
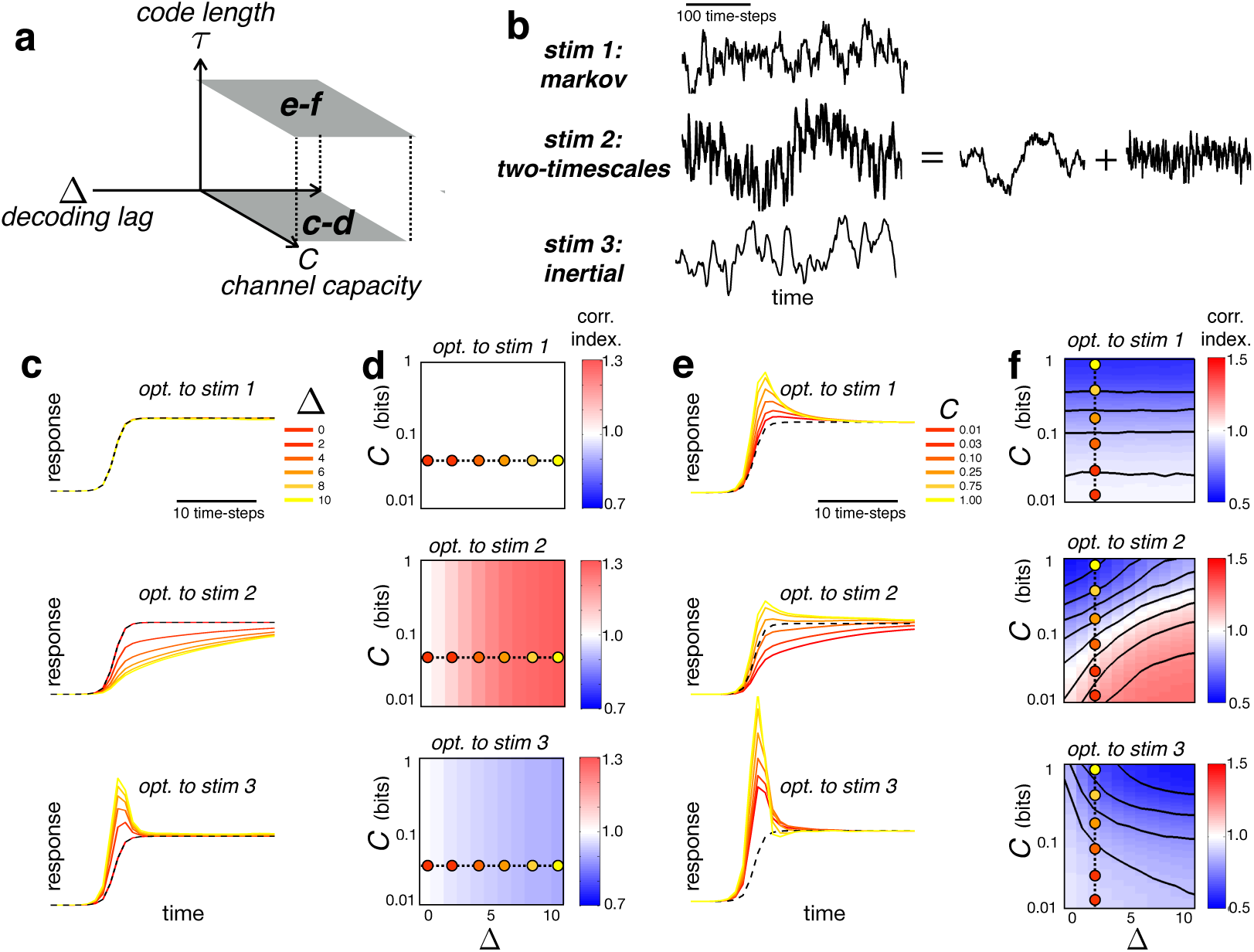
Dependence of optimal code on decoding lag, Δ, code length, *τ*, and channel capacity, *C.* **(a)** We investigated two types of code: instantaneous codes, where *τ* = 0 (panels c-d), and temporally extended codes, where *τ* > 0 (panels e-f). **(b)** Training stimuli used in our simulations. Markov stimulus: the future only depends on the present state. Two-timescale stimulus: sum of two Markov processes that vary over different timescales (shown at right). Inertial stimulus: future depends on stimulus at previous two time-steps. **(c)** Neural responses to probe stimulus (dashed line) after optimising code with varying Δ, and *τ* = 0. Responses are normalised by the final, steady state value. **(d)** Correlation index after optimisation with varying Δ & *C*. Correlation index measures the correlation between neural responses at adjacent timesteps, normalized by the stimulus correlation at adjacent timesteps, i.e., 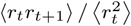 divided by 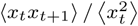. Values greater smaller than 1 indicate that neurons temporally correlate (red) / decorrelate (blue) their input. Filled circles show the parameter values used in panel c. **(e-f)** Same as c-d, but with code optimised for *τ* ≫ 0. Plots in panel e correspond to responses to probe stimulus (dashed line) at varying channel capacity & fixed decoding lag (i.e., Δ = 3, indicated by dashed line in panel f).

With a “two-timescale” stimulus, constructed from two Markov processes that vary over different timescales (Fig 2b, middle panel), the optimal solution was a low-pass filter, to selectively encode the predictive, slowly varying, part of the stimulus. The strength of the low-pass filter increased monotonically with the decoding lag, Δ (Fig 2c-d, middle panels).

Finally, with an “inertial” stimulus, whose future trajectory depended on the previous two states, *y_t_*, and *y*_*t*−1_ (Fig 2b, lower panel), the optimal solution was a high-pass filter, so as to transmit information about velocity. The strength of the high-pass filter also increased monotonically with the decoding lag, Δ (Fig 2c-d, lower panels).

With an instantaneous code, varying the channel capacity, *C*, only rescales responses (relative to the noise amplitude), so as to alter their signal-to-noise ratio. However, the response shape is left unchanged (regardless of the stimulus statistics; Fig 2d). In contrast, with temporally extended codes, where *τ* > 0 (so the stimulus at time *t* + Δ is estimated from the integrated responses between time *t* − *τ* and *t*; Fig 2a) the optimal neural code varies with the channel capacity, *C*. In common with classical efficient coding, at high *C* (i.e. high signal-to-noise ratio) neurons *always* decorrelated their input, regardless of both the stimulus statistics and decoding lag, Δ. Also in common with classical efficient coding, decreasing *C* always led to more correlated responses [7]. However, *unlike* efficient coding, at low to intermediate values of *C* (i.e. intermediate to low signal-to-noise ratio) the optimal code was qualitatively altered by varying the decoding lag, Δ. With the Markov stimulus, increasing Δ had no effect; with the two-timescale stimulus it led to low-pass filtering; and with the inertial stimulus it led to stronger high-pass filtering.

Taken together, “phase diagrams” for optimal, temporally-extended codes show how regimes of decorrelation/whitening (high-pass filtering) and of smoothing (low-pass filtering) are preferred depending on channel capacity, *C*, and decoding lag, Δ. We verified that a qualitatively similar transition from low- to high-pass filtering is also observed with higher dimensional stimuli, and/or more neurons. Importantly, we show that these phase diagrams depend in an essential way on the stimulus statistics already in the linear, gaussian case. We next examined what happens for non-gaussian, high-dimensional stimuli.

### Predictive versus efficient coding of naturalistic stimuli

Natural stimuli exhibit a strongly non-gaussian statistical structure which is essential for human perception [25, 24]. A large body of work has investigated how neurons could efficiently represent such stimuli by encoding their non-redundant, or independent, components [19]. Under fairly general conditions, this is equivalent to finding a sparse code, where each neuron responds selectively to a single, rarely occurring, stimulus feature. For natural images this leads to neurons selective for spatially localised image contours, qualitatively similar to the receptive fields (RFs) of V1 simple cels [8, 26]. For natural movies this leads to neurons selective for a particular motion direction, again similar to observations in area V1 [27].

However, an independent (sparse) temporal code has only been shown to be optimal when: (i) the goal is to maximise information about *past* inputs, i.e, Δ < 0; (ii) at low noise, i.e., at high capacity, *C* ≫ 0. We were interested, therefore, in what happens when these two criteria are violated; for example when neural responses are optimised to encode predictive information, i.e., for Δ ≥ 0.

To explore these questions we modified the objective function of Eq (3) to deal with multi-dimensional stimuli and nongaussian statistics of natural images. To achieve this, we generalized the second term of our objective function to allow optimization of the neural code with respect to higher-order (i.e., beyond covariance) response statistics. Crucially, this modification, described in SI Section 1 and [23], permits—but does not enforce by hand—the sparsity of neural responses. For non-sparse, gaussian stimuli the modification automatically recovers the results of the previous section; for natural stimuli it replicates previous sparse coding results in the limit Δ < 0 and *C* ≫ 0 (see SI Fig 3), without introducing any new tuneable parameters.

We investigated how the optimal neural code for natural stimuli varied with the decoding lag, Δ, while keeping channel capacity, *C*, and code length, *τ*, constant. Stimuli were constructed from 10 × 10 pixel patches drifting stochastically across static natural images (Fig 3a & SI Fig 1; see SI Section 2.2). Neural encoding weights were optimised with two different decoding lags: for Δ = −6 the goal was to efficiently encode the past, while for Δ = 1 the goal was to predict the near future. Figure 3b confirms that the codes indeed are optimal either for efficiency (Δ = −6) or prediction (Δ = 1), as desired.

**Fig. 3.**
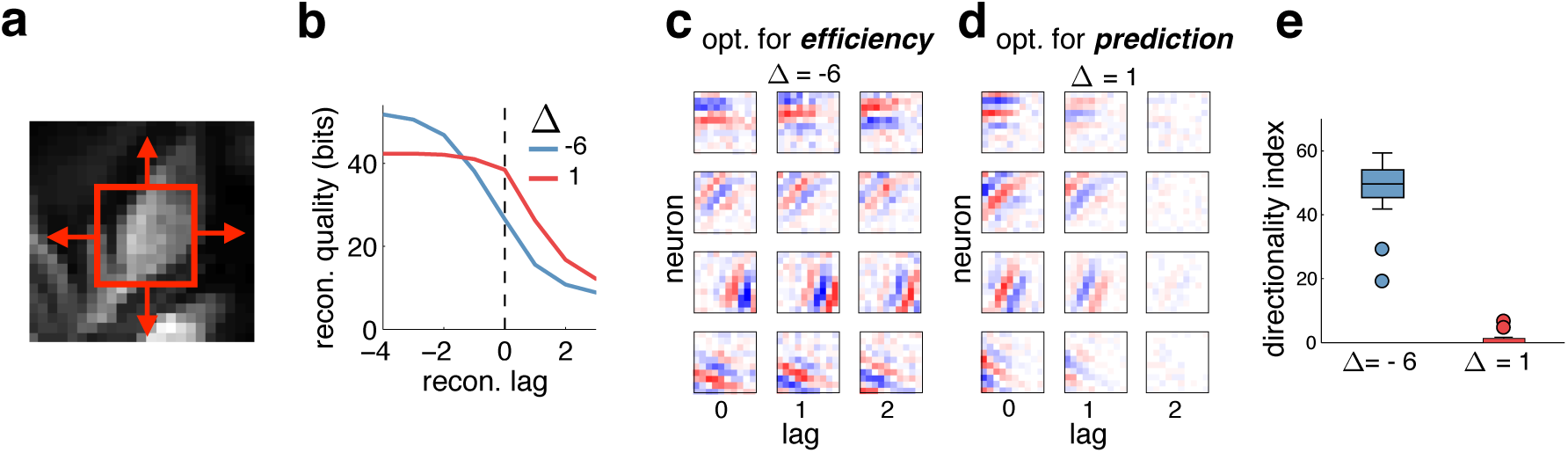
Efficient versus predictive coding of natural stimuli. **(a)** Movies were constructed from a 10×10 pixel patch (red square) which drifted stochastically across static natural images. **(b)** Information encoded by neural responses about the stimulus at varying lag, after optimization with Δ = −6 (blue) and Δ = 2. **(c)** Spatio-temporal encoding filters, for 4 example neurons, after optimisation with Δ = −6. **(d)** Same as panel c, for Δ = 1. **(e)** Directionality index of neural responses, after optimisation with Δ = −6 and Δ = 1. The directionality index measures the percentage change in response to a grating stimulus moving in a neuron’s preferred direction, versus the same stimulus moving in the opposite direction.

After optimisation at both values of Δ, individual neurons were selective to local oriented edge features (Fig 3c-d) [8]. Varying Δ qualitatively altered the temporal features encoded by each neuron, while having little effect on their spatial selectivity. Consistent with previous results in the efficient coding regime [27], single cells at Δ = −6 were responsive to stimuli moving in a preferred direction, as evidenced by spatially displaced encoding filters at different lags (Fig 3c & SI Fig 4a-c), and a high “directionality index” (Fig 3e). In contrast, for predictive coding setup at Δ = 1, cells responded equally to stimuli moving in either direction perpendicular to their encoded stimulus orientation. This was evidenced by spatio-temporally separable receptive fields (SI Fig 4d-f) and directionality indexes near zero. This qualitative difference between the efficient and predictive code for natural movies was highly surprising, and we sought to understand its origins.

### Trade-off between sparsity and predictive power

To gain an intuitive understanding of how the optimal code varies with decoding lag Δ, we constructed artificial stimuli from overlapping “gaussian bumps” which drifted stochastically along a single spatial dimension (Fig 4a; SI Section 2.3). While simple, this stimulus captured two key aspects of the natural movies: first, the gaussian bumps drifted smoothly in space, resembling the stochastic global motion over the image patches; second, the stimulus also had a sparse latent structure.

**Fig. 4.**
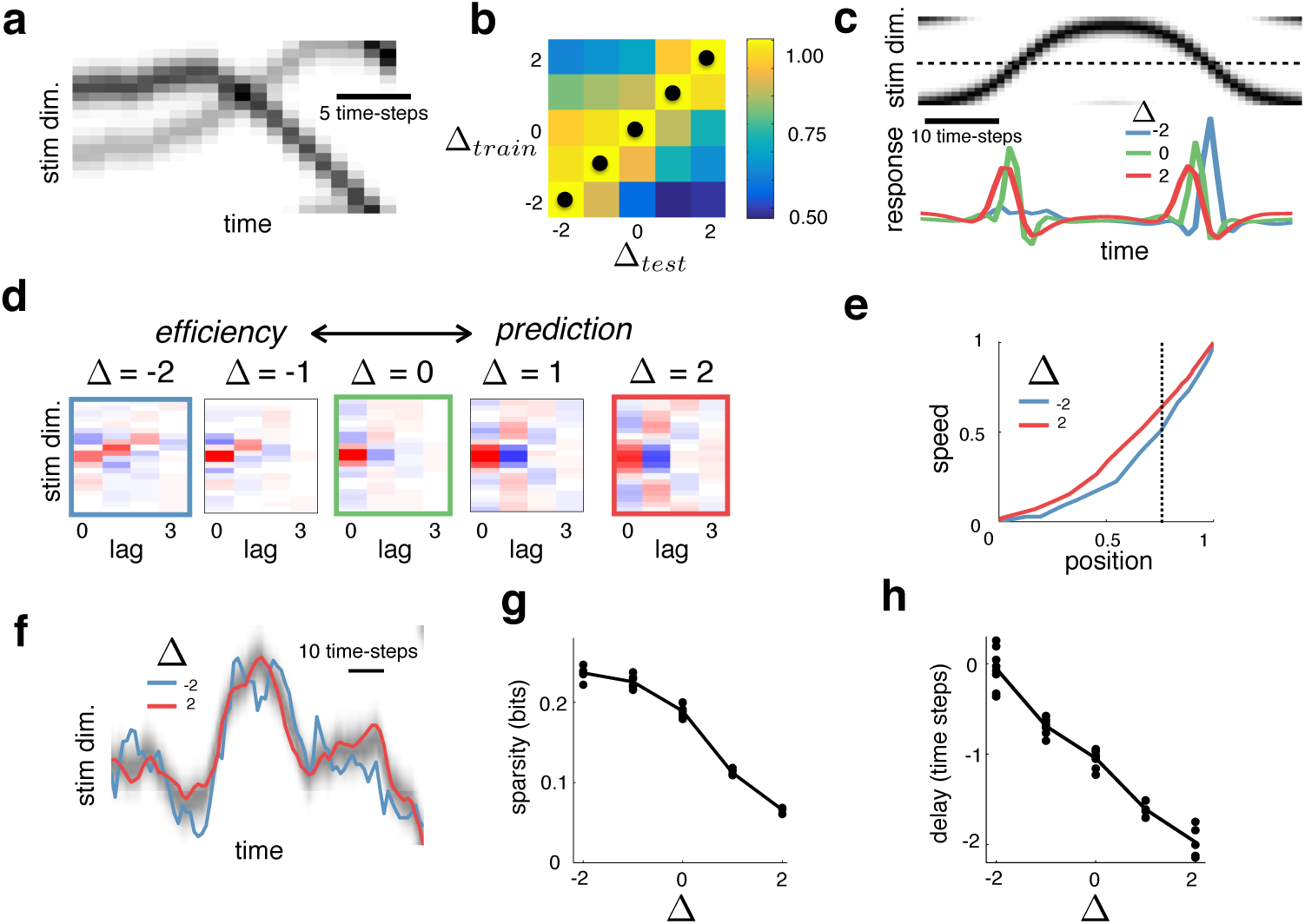
Efficient versus predictive coding of a “gaussian-bump” stimulus. **(a)** Stimuli consisted of gaussian-bumps that drifted stochastically along a single spatial dimension (with circular boundary conditions). **(b)** Information encoded by neural responses about the stimulus at varying lag, Δ_test_, after optimization with varying Δ_train_. Black dots indicate the maximum for each column. **(c)** Response of example neuron to a test stimulus (above), after optimisation with Δ = −2 (blue), Δ = 0 (green), and Δ = 2 (red). **(d)** Spatio-temporal encoding filters for an example neuron, after optimisation with different Δ. **(e)** Circular correlation between the reconstructed speed of a moving gaussian blob and its true speed, versus the circular correlation between the reconstructed position and its true position, obtained from neural responses optimised with Δ = ±2 (red and blue curves). Curves were obtained by varying *γ* in Eq (3), to find codes with different channel capacities. **(f)** Linear reconstruction of the stimulus trajectory, obtained from neural responses optimised with Δ = ±2 (red and blue curves). The full stimulus is shown in grayscale. While coding capacity was chosen to equalize the mean reconstruction error for both models (vertical dashed line in panel e), the reconstructed trajectory was much smoother for the predictive (red) than for the efficient (blue) coding model. **(g)** Response sparsity (defined as the negentropy of neural responses), versus Δ (dots = individual neurons; line = population average). **(h)** Delay between stimulus presented at a neuron’s preferred location and each neuron’s maximum response, versus Δ.

We optimised the neural code with Δ ranging from −2 to 2, holding the channel capacity, *C*, and code length, *τ*, constant. Fig 4b confirms that highest performance was achieved when the reconstruction performance was evaluated at the same lag for which each model was trained. This simpler setup recapitulated the surprising result we obtained with naturalistic stimuli: namely, that when Δ < 0 neurons were selective to a single preferred motion direction, while when ≥ > 0 neurons responded equally to stimuli moving from either direction into their receptive field (Fig 4c-d).

Predicting the future state of the stimulus requires estimating its current motion direction and speed. How is it possible, then, that an optimal predictive code (Δ > 0) results in neurons being unselective to motion direction? This paradox is resolved by realising that it is the information encoded by the *entire neural population* that counts, not the information encoded by individual neurons. Indeed, when we looked at the information encoded by the neural population, we did find what we had originally expected: when optimised with Δ > 0, the neural population as a whole encoded significantly more information about the stimulus velocity than its position (relative to when Δ < 0), despite the fact that individual neurons were unselective to motion direction (Fig 4e-f).

The change in coding strategy that is observed as one goes from efficient (Δ < 0) to predictive coding (Δ ≥ 0) is in part due to a tradeoff between cells maintaining sparse responses (which is efficient) and responding quickly to stimuli within their RF (which helps predictions). Intuitively, to be efficient and respond with greatest selectivity, the neuron first has to wait to process and recognize the “complete” stimulus feature; unavoidably, however, this entails a processing delay and leaves no information to be encoded predictively. This can be seen in Fig 4g-h, which shows how both the response sparsity and delay to stimuli within a cell’s RF decrease with Δ. In SI section 3.4 we describe in detail why this trade-off between efficiency and prediction leads to direction selective filters when Δ < 0, but not when Δ > 0 (SI fig. 5).

Beyond the effects on the optimal code of various factors explored in detail in the main paper, our framework further generalises previous efficient and sparse coding results to factors listed in Table 1 and discussed in SI Section 3.5. For example, decreasing the capacity, *C* (while holding Δ constant at −2) resulted in neurons being unselective to stimulus motion (SI fig 6a), with a similar result observed for increased input noise (SI fig 6b). Thus, far from being generic, traditional sparse temporal coding, in which neurons responded to local motion, was only observed in a specific regime (i.e., Δ < 0, *C* ≫ 0 and low input noise, *n* ~ 0).

## Discussion

Efficient coding has long been considered a central principle for understanding early sensory representations [4, 5], with well-understood implications and generalizations [9, 28]. It has been successful in predicting many aspects of neural responses in early sensory areas directly from the low-order statistics of natural stimuli [7, 29, 30, 31, 24], and has even been extended to higher-order statistics and central processing [32, 33]. However, a criticism of the theory is that it implicitly treats all sensory information as equal, despite empirical evidence that neural systems prioritise behaviourally relevant (and not just statistically likely) stimuli [34]. To overcome this limitation, Bialek and colleagues proposed an alternative theory, called predictive coding, which posits that neural systems encode maximal information about future inputs, given fixed information about the past [11, 12]. This theory is motivated by the fact that stimuli are only useful for performing actions when they are predictive about the future.

Compared to efficient coding, predictive coding has remained relatively unexplored (though see later for alternative definitions of predictive coding, which have recieved more attention). Existing work only considered the highly restrictive scenario where neurons maximise information encoded in their *instantaneous* responses [12, 21, 17]. In this case (and subject to some additional assumptions, such as gaussian stimulus statistics and instantaneous encoding filters), predictive coding is formally equivalent to slow feature analysis [18]. This is the exact opposite of efficient coding, which (at low noise/high capacity) predicts that neurons should temporally decorrelate their inputs [14].

To clarify the relation between efficient and predictive coding, we developed a unified framework that can treat both theories [22, 11, 23]. We investigated what happens when the neural code is optimised to be both predictive and temporally efficient (Fig b). In this case, the optimal code depends critically on the channel capacity (i.e. signal-to-noise ratio), which describes how much information the neurons can encode about their input. At high capacity (i.e. low-noise), neurons always temporally decorrelate their input. At finite capacity (i.e. mid- to high-noise), however, the optimal neural code varies qualitatively depending on whether the goal is to reliably predict the future or efficiently reconstruct the past.

When we investigated predictive coding of natural stimuli, we found solutions that are qualitatively different from known sparse coding results, in which individual neurons are tuned to directional motion of local edge features [27]. In contrast, we found that neurons optimised for predictive coding are selective for motion speed but not direction (Fig 3 and SI Fig 4). Surprisingly, however, the neural population as a whole encodes motion even more accurately than before (Fig 4e). We show that these changes are due to an implicit trade-off between maintaining a sparse code (which is efficient) and responding quickly to stimuli within each cell’s RF (which aids predictions; Fig 4g-h).

It is notable that, in our simulations, strikingly different conclusions are reached by analysing single neuron responses versus, the responses of the entire neural population. Specifically, looking only at single neurons responses would lead one to conclude that when performing predictive coding, neurons *did not* encode motion direction; looking at the neural population responses reveals that the opposite is true. This illustrates the importance of population-level analyses of neural data, and how, in many cases, single neuron responses can give a false impression of which information is represented by the population.

A major challenge in sensory neuroscience is to derive the observed cell-type diversity in sensory areas from a normative theory. For example, in visual area V1, one observes a range of different cell-types, some of which are have spatio-temporally seperable RFs, and others which do not [35, 36]. The question arises, therefore, whether the difference between cell-types emerges because different subnetworks fulfill qualitatively different functional goals. One hypothesis, suggested by our work, is that cells with seperable RFs have evolved to encode predictive information, while cells with non-seperable RFs evolved to optimise coding efficiency. More generally, the same hypothesis could explain the existence of multiple cell-types in the mammalian retina [37], with each cell-type mosaic implementing an optimal code for a particular choice of optimisation parameters, e.g., channel capacity or prediction lag.

Testing such hypotheses rigorously against quantitative data would require us to generalise our work to nonlinear encoding and decoding models (Table 1, final row). Here we focused on a linear decoder to lay a solid theoretical foundation and permit direct comparison with previous sparse coding models, which also assumed a linear decoder [27, 8, 26]. In addition, a linear decoder forces our algorithm to find a neural code for which information can be easily extracted by downstream neurons performing biologically plausible operations. While the linearity assumptions simplify our analysis, the framework can easily accommodate non-linear encoding and decoding. For example, we previously used a “kernel” encoding model, where neural responses are described by a non-parametric & non-linear function of the input [23]. Others have similarly used a deep convolutional neural network as an encoder [38].

In the future it would be interesting to investigate how our ideas relate to sensory processing in a hierarchy. Hierarchical processing has long been discussed in the context of efficient coding [24], where neurons at each layer are assumed to remove residual statistical dependencies in their inputs [39, 19]. In contrast, hierarchical predictive coding, in which neurons at each layer encode maximal information about their future inputs, has not yet been explored.

“Predictive coding” has been used to describe different approaches. Here, we understood the term in information-theoretic context, implying that neurons preferentially encode stimuli that carry information about the future [11]. However, predictive coding has also been used to imply that neurons encode “surprising” stimuli, i.e., those not predictable from past inputs [4, 40, 41]. Elsewhere, predictive coding describes a particular type of hierarchical processing, in which feed-forward projections encode an error signal, equal to the difference between bottom-up sensory inputs and top-down predictions from higher sensory areas [42, 43]. These alternative definitions of predictive coding are not equivalent. For example, sensory stimuli can be surprising based on past inputs, but not predictive about the future [44]. Likewise, previous theories of hierarchical predictive coding do not address which sensory information should be preferentially encoded or alternatively, discarded. Clarifying the relationship between these inequivalent definitions of predictive coding and linking them mathematically to coding efficiency provided one of the initial motivations for our work. In past work, alternative coding theories are often expressed using very different mathematical frameworks, impeding comparison between them, and sometimes leading to confusion. In contrast, by using a single mathematical framework to compare different theories— efficient, sparse and predictive coding—we were able see exactly how they relate to each other, the circumstances under which they make opposing or similar predictions, and what happens when they are combined.

1 The term ‘predictive coding’ has been used previously to describe several different approaches. In our work, we use the definition given by [11], were nrurons encode maximal information about the future, given information encoded about the past. Alternative definitions are described in the discussion.

2 In other words, previous efficient coding models maximised the encoded information *rate* at time *t*, while previous predictive coding models maximised the total encoded information at time *t.*

## Acknowledgements

This work was supported in part by the Austrian Science Fund grant FWF P25651.

